# Inhibition of glycolysis in tuberculosis-mediated metabolic rewiring reduces HIV-1 spread across macrophages

**DOI:** 10.1101/2024.08.14.607908

**Authors:** Zoï Vahlas, Natacha Faivre, Sarah C. Monard, Quentin Hertel, Mariano Maio, Joaquina Barros, Alexandre Lucas, Thien-Phong Vu Manh, Myriam Ben Neji, Marcelo Corti, Renaud Poincloux, Fabien Blanchet, Brigitte Raynaud-Messina, Fabien Letisse, Olivier Neyrolles, Geanncarlo Lugo-Villarino, Luciana Balboa, Christel Vérollet

## Abstract

Tuberculosis (TB) is a significant aggravating factor in individuals living with human immunodeficiency virus type 1 (HIV-1), the causative agent for acquired immunodeficiency syndrome (AIDS). Both *Mycobacterium tuberculosis* (Mtb), the bacterium responsible for TB, and HIV-1 target macrophages. Understanding how Mtb subverts these cells may facilitate the identification of new druggable targets. Here, we explored how TB can induce macrophages to form tunneling nanotubes (TNT), promoting HIV-1 spread. We found that TB triggers metabolic rewiring of macrophages, increasing their glycolytic ATP production. Using pharmacological inhibitors and glucose deprivation, we discovered that disrupting aerobic glycolysis significantly reduces HIV-1 exacerbation in these macrophages. Glycolysis is essential for tunneling nanotubes (TNT) formation, which facilitates viral transfer and cell-to-cell fusion and induces the expression of the sialoadhesin Siglec-1, enhancing both HIV-1 binding and TNT stabilization. Glycolysis did not exacerbate HIV-1 infection when TNT formation was pharmacologically prevented, indicating that higher metabolic activity is not sufficient per se to make macrophages more susceptible to HIV-1. Overall, these data might facilitate the development of targeted therapies aimed at inhibiting glycolytic activity in TB-induced immunomodulatory macrophages to ultimately halt HIV-1 dissemination in co-infected patients.

## INTRODUCTION

Tuberculosis (TB) and acquired immunodeficiency syndrome (AIDS) are among the deadliest diseases caused by single infectious agents. A significant issue in the AIDS epidemic is the synergy between the human immunodeficiency virus type 1 (HIV-1) and *Mycobacterium tuberculosis* (Mtb), the etiological agents of AIDS and TB, respectively. Globally, Mtb is the most frequent co-infection in patients with HIV-1 and represents a major risk factor for increased morbidity and mortality (WHO Global tuberculosis report 2021). Addressing the TB challenge within the AIDS epidemic requires a comprehensive understanding of the pathophysiology of HIV/Mtb co-infection, including the role of immunometabolism (Esmail et al., 2018).

Despite CD4^+^ T lymphocytes being the primary target for HIV-1, macrophages provide a crucial niche for both pathogens, allowing Mtb to evade immune responses and HIV-1 to persist and replicate, thereby exacerbating the impact of co-infection (Cohen et al., 2018; Mayer-Barber and Barber, 2015; Tan and Russell, 2015). On the one hand, lung macrophages are the main cell target for Mtb because they are the primary immune cells responsible for engulfing and attempting to eliminate pathogens through phagocytosis, providing an intracellular environment that the bacteria can exploit to survive and replicate. On the other hand, infected macrophages are found in tissues of HIV^+^ patients and simian immunodeficiency virus (SIV)-infected non-human primates (NHPs), playing an important role in the pathogenesis (Honeycutt et al., 2017; Honeycutt et al., 2016; Rodrigues et al., 2017). In addition, multiple studies demonstrate that tissue macrophages, such as microglia as well as urethral, gut and lung macrophages, can be reservoirs for HIV-1 in patients undergoing antiretroviral therapy (Ganor et al., 2019; Ganor et al., 2013; Sattentau and Stevenson, 2016). In the lung, for example, the macrophage compartment is the main target of HIV-1 (Avalos et al., 2016; Jambo et al., 2014; Schiff et al., 2021). Our team showed recently that the abundance of lung macrophages becomes augmented in NHPs with active TB and exacerbated in those co-infected with SIV, acquiring an immunomodulatory phenotype distinguished by the overactivation of the interleukin-10 (IL-10)/signal transducer and activator of transcription 3 (STAT3) axis (Dupont et al., 2022; Dupont et al., 2020; Souriant et al., 2019). This phenotype is closely related to the so called “M(IL-10)” activation program (Murray et al., 2014), found abundantly in the pleural cavity of patients with active TB, and reproduced *in vitro* by exposure of human monocytes to TB-associated microenvironments (Lastrucci et al., 2015). Further work with these TB-induced immunomodulatory macrophages demonstrated an increased susceptibility to HIV-1 replication and spread *via* the formation of tunneling nanotubes (TNTs), which facilitate the transfer of the virus between macrophages, trigger their fusion, leading to the formation of highly virus-productive multinucleated giant cells (MGCs) (Souriant et al., 2019). Of note, MGC are considered the pathological hallmarks of HIV-1 infection in macrophages (Orenstein, 2000; Verollet et al., 2015; Verollet et al., 2010). TNTs are membranous channels containing F-actin and connecting cells together over long distances, and they can be hijacked by pathogen to circumvent the immune system (Dupont et al., 2018; Zurzolo, 2021). However, the molecular mechanisms linking TB infection and TNT formation, stability and function in immunomodulatory macrophages are not well understood, and uncovering these might contribute to the development of novel targeted therapies.

Chronic host-pathogen interactions in TB result in extensive metabolic remodeling in both the host and the pathogen (Huang et al., 2019; Kumar et al., 2019; Llibre et al., 2021). In fact, the success of Mtb as a pathogen largely relies on its ability to adapt to the intracellular milieu of macrophages and use of their metabolic activity to its advantage. In chronic infectious diseases, there is often a shift in the macrophage activation program from the M1 phenotype, which primarily relies on aerobic glycolysis, towards the M2 phenotype, which depends heavily on oxidative phosphorylation (OXPHOS), at the site of inflammation (Biswas and Mantovani, 2012; Lugo-Villarino et al., 2011; Saha et al., 2017). This metabolic rewiring is associated with the adaptive immune transition from acute to chronic phases (Cumming et al., 2018; Gleeson et al., 2016; Marin Franco et al., 2020; O’Maoldomhnaigh et al., 2021; Vrieling et al., 2020). In general, macrophages infected by live Mtb acquire the M1 phenotype, characterized by elevated production of pro-inflammatory molecules. They rely on aerobic glycolysis and the pentose phosphate pathway to meet their bioenergetic and metabolic requirements. However, results can vary based on differences in Mtb strains, multiplicity of infection, macrophage origins, and measurement timepoints (Cumming et al., 2018; Gleeson et al., 2016; Marin Franco et al., 2020; Vrieling et al., 2020). In such M1 macrophages, OXPHOS and fatty acid oxidation are dampened. Nonetheless, the untimely overproduction of lactate, the end-product of aerobic glycolysis, disrupts macrophage metabolism, leading to an attenuated glycolytic shift upon subsequent stimulation with irradiated Mtb and reduced pro-inflammatory cytokine production (O’Maoldomhnaigh et al., 2021). Also, a shift from glycolysis to OXPHOS is observed in M1 macrophages when they are exposed to the acellular fraction of pleural effusions (PE) from TB patients (TB-PE), considered hereafter as a genuine TB-associated microenvironment (Marin Franco et al., 2020). Of note, human monocyte differentiation under TB-PE yields immunoregulatory macrophages (Lastrucci et al., 2015). Thus, different metabolic pathways are prevalent in macrophages depending on their ontogeny, the state of TB (active or latent) and the virulence of the pathogen (live or irradiated Mtb); these states can be remarkably reversible depending on environmental cues (Beste et al., 2013; Cano-Muniz et al., 2018; Cumming et al., 2018; de Carvalho et al., 2010; Llibre et al., 2021; Pandey and Sassetti, 2008; Zimmermann et al., 2017).

While the role of macrophage metabolism is well described in TB, little is known for HIV-1 host cells (Saez-Cirion and Sereti, 2021; Shehata et al., 2017). Our knowledge about how metabolism affects HIV-1 infection comes essentially from studies in CD4 T lymphocytes (Clerc et al., 2019; Craveiro et al., 2013; Loisel-Meyer et al., 2012; Saez-Cirion and Sereti, 2021; Valle-Casuso et al., 2019). To be efficiently infected by HIV-1, these cells must be metabolically active. In this regard, the increase in both aerobic glycolysis and OXPHOS is crucial for the early step of HIV-1 infection (Clerc et al., 2019; Valle-Casuso et al., 2019). In the case of macrophages, it was shown that HIV-1 modifies their metabolic status (Castellano et al., 2019; Hollenbaugh et al., 2011; Saez-Cirion and Sereti, 2021). A recent report indicated that a metabolic shift toward aerobic glycolysis can reactivate HIV-1 replication in macrophages (Real et al., 2022). However, the underlying mechanisms and consequences remain unknown. While the study of immunometabolism on HIV-1 infection and progression is an emerging field, very little research has focused on macrophage metabolism, especially in the context of coinfection with Mtb.

In summary, our research demonstrates that TB significantly exacerbates HIV-1 infection by inducing metabolic reprogramming in macrophages, leading to increased glycolytic ATP production. This metabolic shift promotes the formation of TNTs, which facilitate viral transfer and cell-to-cell fusion. The study highlights the critical role of glycolysis in TNT formation and the subsequent expression of the sialoadhesin Siglec-1, which enhances HIV-1 binding and TNT stabilization. Importantly, inhibiting glycolysis significantly reduces HIV-1 exacerbation in TB-infected macrophages, suggesting that targeting glycolytic pathways could be a promising therapeutic strategy to prevent HIV-1 dissemination in co-infected patients.

## RESULTS

Human monocytes differentiate into highly glycolytic immunomodulatory macrophages upon exposure to TB-associated microenvironment Understanding the TB-associated microenvironment in the context of HIV-1 co-infection is essential because TB significantly worsens the prognosis for individuals infected with HIV-1. Although it is rare to find cells co-infected with both pathogens, studying these microenvironments is vital since TB and HIV-1 often occupy adjacent zones, such as in the lungs, leading to enhanced viral dissemination and immune system manipulation. To determine whether the metabolic profile of macrophages is modulated in TB-associated microenvironments, we employed a previously described *in vitro* model (Dupont et al., 2022; Dupont et al., 2020; Lastrucci et al., 2015; Souriant et al., 2019), which consists of differentiating primary human monocytes into macrophages in the presence of a TB-associated microenvironment, such as conditioned medium generated from Mtb-infected macrophages (cmMTB). These macrophages display an immunomodulatory phenotype characterized by a CD16^+^CD163^+^MerTK^+^PD-L1^+^ receptor signature, nuclear translocation of phosphorylated STAT3, and functional properties like protease-dependent motility, suppression of T cell activation, and susceptibility to Mtb or HIV-1 infection (Dupont et al., 2022; Dupont et al., 2020; Lastrucci et al., 2015; Souriant et al., 2019). Conditioned medium of non-infected macrophages (cmCTR) was used as control. We initially focused on the two main pathways that produce cell energy in the form of adenosine triphosphate (ATP), which are aerobic glycolysis and OXPHOS. The intracellular rate of ATP production was evaluated by the Seahorse technology at day three of monocyte differentiation into macrophages with either cmCTR or cmMTB. First, we found that total ATP production was strongly increased in cmMTB-differentiated macrophages compared to control cells (**Figure 1A**). The measurements of basal extracellular acidification rate (ECAR) and basal oxygen consumption rate (OCR) were used to calculate ATP production rate from glycolysis (GlycoATP) and mitochondrial OXPHOS (MitoATP), respectively, at different time points (**Figure 1 and S1A-B**). We found that the intracellular GlycoATP production was significantly increased in cmMTB-differentiated macrophages compared to control cells (**Figure 1B**). Kinetic analyses revealed that the increase in GlycoATP production began as early as day one and peaked at day three of monocyte differentiation under cmMTB (**Figure S1A**). However, this change in the metabolic state was transitory, as the difference in GlycoATP production observed between cmCTR and cmMTB conditions vanished by day six (**Figure S1A**).

**Figure 1.**
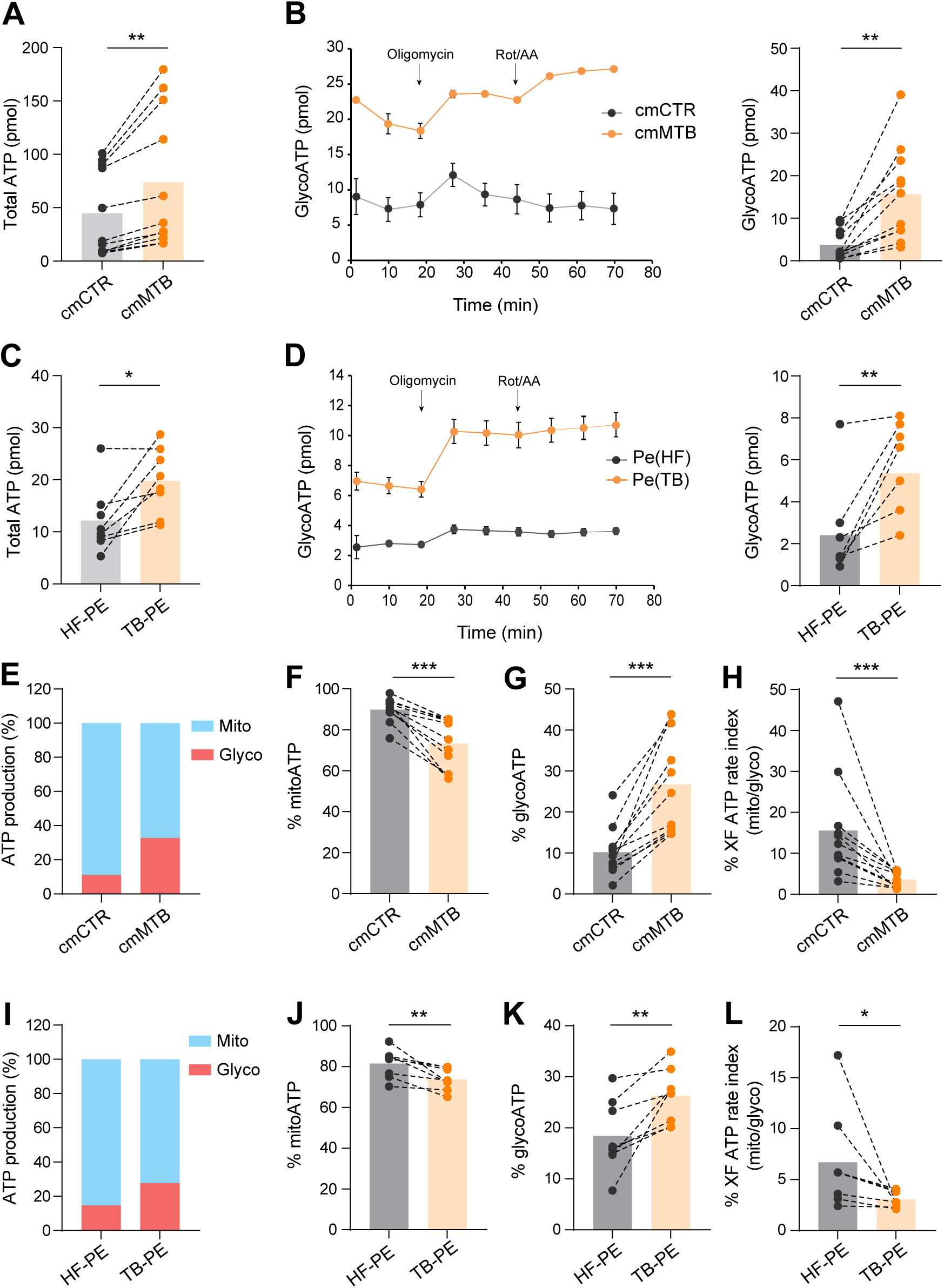
TB-associated microenvironments increase aerobic glycolysis in immunoregulatory macrophages. Monocytes from healthy subjects were treated either with conditioned medium from mock-(cmCTR, grey) or Mtb-infected macrophages (cmMTB, orange), or with heart failure (HF-PE, grey) or pleural effusions (PE) from TB patients (TB-PE, orange) for three days and analyzed using an Agilent Seahorse XFe24 Analyzer. (**A, C**) Dot plots showing total ATP production. (**B, D**) GlycoATP rate after addition of Oligomycin and Rotenone/Antimycin A (ROT/AA) over time (left) and dot plots showing total GlycoATP production (right). (**E, I**) Percentages of MitoATP and GlycoATP production relative to overall ATP production, representative experiments. (**F, J**) Dot plots showing the percentages of MitoATP. (**G, K**) Dot plots showing the percentages of GlycoATP. (**H, L**) Scattered plots showing ATP rate index (% XF). Each circle within vertical plots represents a single donor. Histograms represent mean values. Statistical analysis: t-test data with normal distribution; *, p≤0.05; **, p≤0.01; ***, p≤0.001.

Importantly, high ATP production, and especially GlycoATP production, was also observed in monocyte differentiation into immunomodulatory macrophages under another TB-associated microenvironment, such as the acellular fraction of TB-PE. This contrasted with conditioning with PE obtained from patients with heart failures (HF-PE), which was used as a control (**Figure 1C-D**). Simultaneously, the level of MitoATP also increased in cmMTB- or TB-PE-differentiated macrophages compared to their control counterparts (**Figure S1B-C**), displaying similar kinetics to GlycoATP production. Also, the maximal respiration and spare respiratory capacity were slightly increased in cmMTB-differentiated macrophages relative to those conditioned with cmCTR (**Figure S1D-E**), as measured by the Seahorse Mito Stress^TM^ assay. Since MitoATP is different between cmMTB-and cmCTR-differentiating macrophages, the number of mitochondria was assessed using transmission electron microscopy (TEM) (**Figure S1F-G**), and mitochondrial biomass was measured by flow cytometry (**Figure S1H**); both parameters were found to be comparable between the two conditions. Using the MitoSOX^TM^ fluorescent probes to measure intracellular superoxide formation, no statistical difference in oxidative stress was observed between these two cell populations (**Figure S1I**). This suggests that the increase in maximal respiration and spare respiratory capacity are not a consequence of cellular stress. Next, to understand the relative contributions of mitochondrial respiration and glycolysis to the bioenergetic profile of macrophages differentiated under TB-associated microenvironments, we compared the relative utilization of mitochondrial *versus* glycolytic pathway for ATP production between cmMTB- and cmCTR-differentiated macrophages at day three. Approximately 90% of ATP production in macrophages differentiated with cmCTR came from OXPHOS; this parameter was reduced to 70% when conditioned with cmMTB (**Figure 1E-F**). Consistently, the percentage of GlycoATP increased from 10% to more than 25% in cmMTB-differentiated macrophages compared to control cells (**Figure 1E and G**), leading to an overall decrease in the Mito/Glyco ATP ratio (**Figure 1H**). Similar results were obtained with the use of TB-PE (**Figure 1I-L**), demonstrating an increased use of the glycolytic pathway in immunomodulatory macrophages differentiated under different TB-associated microenvironments.

The shift to aerobic glycolysis by TB was further supported by a significant enrichment of glycolytic genes observed in cmMTB-differentiated macrophages compared to control cells (**Figure 2A**). This finding was obtained through the re-analysis of our previously published genome-wide transcriptomic data (GEO submission GSE139511) using a GSEA-based approach (Dupont et al., 2022). By contrast, genes of the OXPHOS pathway failed to be enriched in the cmMTB-differentiated macrophages (**Figure S1J**). To validate this *in silico* analysis, glucose uptake was measured by fluorescent d-glucose analog 2-[N-(7-nitrobenz-2-oxa-1,3-diazol-4-yl) amino]-2-deoxy-D-glucose (2-NBDG). The results indicate a significantly elevated uptake in cmMTB-differentiated macrophages compared to control cells (**Figure 2B**). This increase was reflected in the higher glucose consumption found in the extracellular medium of these macrophages, as measured dynamically using a nuclear magnetic resonance (NMR)-based metabolomic approach (**Figure 2C**). This approach also revealed that the production of extracellular lactate by cmMTB-differentiated macrophages was enhanced compared to controls (**Figure 2D**). The elevated levels of lactate release were also verified using classical spectrophotometric analysis in macrophages differentiated under cmMTB or TB-PE conditions, compared to those with cmCTR or HF-PE, respectively (**Figure 2E-F**). Glycolytic reprogramming taking place in macrophages is orchestrated by the hypoxia-inducible factor-1 alpha (HIF-1α), which increases the expression of glycolytic enzymes and pro-inflammatory cytokines (Gleeson and Sheedy, 2016; Palsson-McDermott et al., 2015). Thus, HIF-1α expression was examined in our experimental conditions. Accordingly, while the protein expression level of HIF-1α was not modified between the two cell populations (**Figure 2G**), only cmMTB-differentiated macrophages exhibited its frequent translocation from the cytoplasm to the nucleus (**Figure 2H-I**), suggesting that TB-associated microenvironments favor glycolysis through HIF-1α activation.

**Figure 2.**
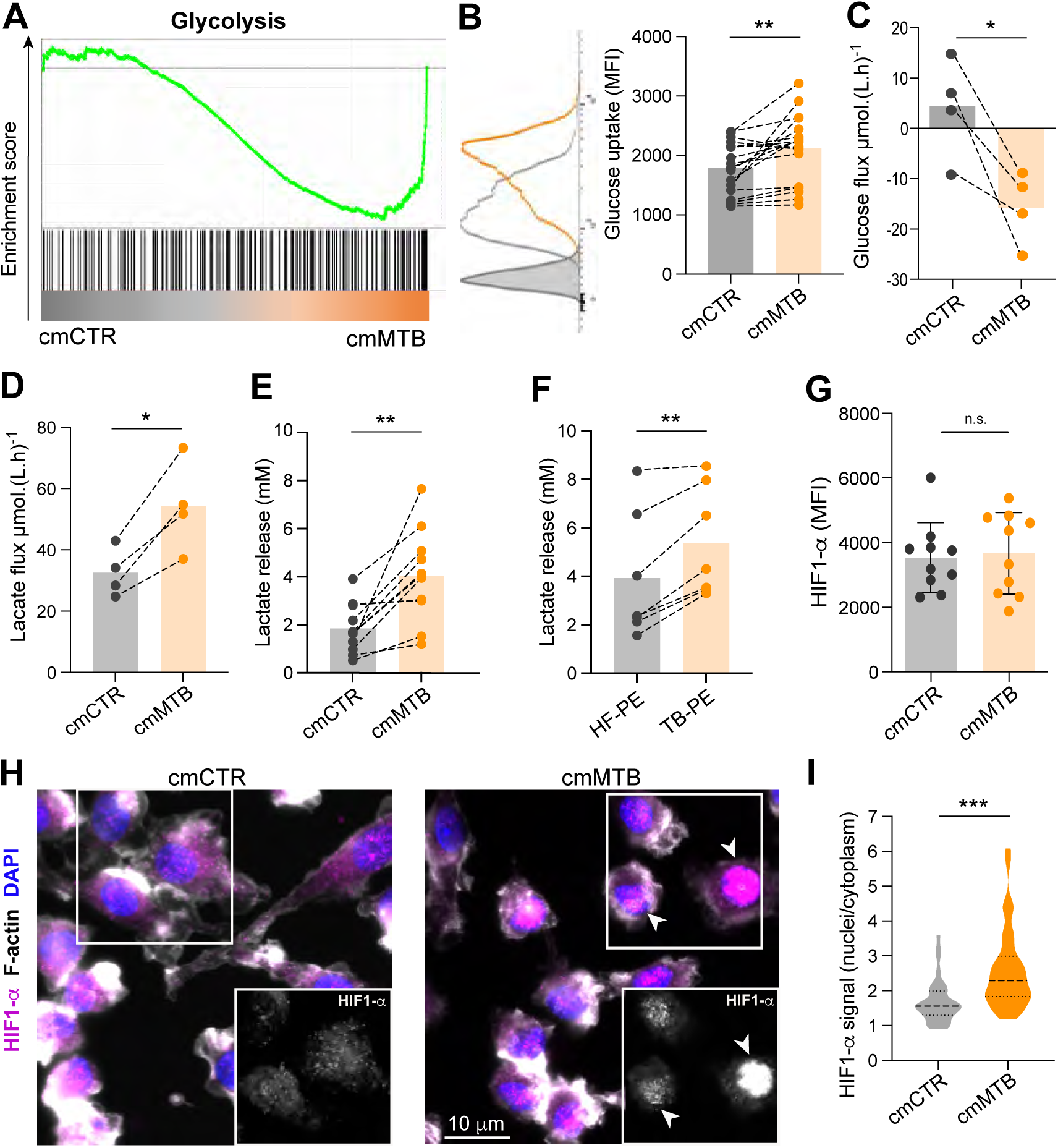
TB-associated microenvironments and Mtb infection increases aerobic glycolysis in immunoregulatory macrophages. (**A-I**) Monocytes were treated either with conditioned medium from mock- (cmCTR, grey) or Mtb-infected macrophages (cmMTB, orange), or with heart failure (HF-PE, grey) or pleural effusions from TB patients (TB-PE, orange). (**A**) Gene set enrichment plot of the glycolytic genes (hallmark collection of MSigDB). This plot shows the distribution of the glycolysis gene set between macrophages exposed to cmCTR versus cmMTB for three days. The skewing of the genes to the right and the negative normalized enrichment score (NES = −1.66) indicate enrichment of genes related to glycolysis in macrophages exposed to cmMTB versus cmCTR. (FDR = 0.005). (**B**) Glucose uptake was assessed by flow cytometry using 2-NBDG (2-(7-Nitro-2,1,3-benzoxadiazol-4-yl)-D-glucosamine) staining in cells treated with cmCTR versus cmMTB for three days. Representative histograms (left) and dot plots showing the geomean fluorescence intensity (MFI). (**C-D**) Supernatant of cmCTR and cmMTB-treated cells was collected at different time points post-treatment and glucose (D) and lactate (E) flux (mmol.(L.h)^-1^) was assessed by RMN analysis for n= four donors. (**E-F**) Dot plots showing the concentration (mM) of lactate in the supernatants at day three. (**G**) HIF1-α expression was assessed by flow cytometry at day three. Mean +/- SD is shown. (B-G) Each circle represents a single donor. Histograms represent mean values. (**H-I**) HIF-1α localization was assessed by IF in macrophages exposed to cmMTB versus cmCTR. (F) Representative immunofluorescence images: HIF-1α (magenta), F-actin (phalloidin, grey) and nuclei (DAPI, blue). Scale bar, ten µm. White arrowheads show HIF-1α translocation in the nucleus. (I) Quantification of the ratio of HIF-1α signal intensity in the nucleus versus the cytoplasm. 20 cells/conditions, n= three donors. Statistical analysis: t-test data with normal distribution; *, p≤0.05; **, p≤0.01; ***, p≤0.001; n.s not significant

### Glycolysis induced by TB-associated microenvironments exacerbates HIV-1 infection of immunomodulatory macrophages

Since our immunomodulatory macrophage model is permissive to HIV-1 infection (Dupont et al., 2022; Dupont et al., 2020; Lastrucci et al., 2015; Souriant et al., 2019), we investigated whether its heightened glycolytic activity contributes to this susceptibility. To achieve this, the metabolism of cmMTB-differentiated macrophages was modulated using a pharmacological approach: UK5099 was used to partially block the entry of pyruvate into the mitochondria, thereby enhancing glycolysis. Additionally, oxamate and GSK 2837808A were employed to specifically target lactate dehydrogenase, the enzyme responsible for converting pyruvate to lactate, thus diminishing glycolysis. (**Figure S2A**). CmMTB-differentiated macrophages were incubated with the indicated drugs for 24 h before HIV-1 infection (**Figure S2B**). A VSVG pseudotyped-NLAD8 strain was used to enhance HIV-1 entry regardless of the presence of CD4/CCR5 entry receptors that might be affected by metabolic changes. First, the multiple drug treatments did not affect cell viability, as verified by flow cytometry (**Figure S2C**) and immunofluorescence (**Figure 3A and S2D**). The effect of the drugs on glycolysis was verified by the modulation of lactate release. As expected, diminished glycolysis reduced the release of lactate by cmMTB-differentiated macrophages, while enhanced glycolysis slightly increased it (**Figure 3B and S2E**). Importantly, enhanced glycolysis significantly augmented the number of HIV-1-infected cmMTB-differentiated macrophages, as evidenced by an augmented intracellular detection of HIV-1-Gag protein expression by immunofluorescence (**Figure 3A and 3C**). In contrast, the HIV-1 infection was reduced by more than two-fold under when glycolysis was diminished (**Figure 3C and S2F**). These contrasting effects in HIV-1 infection through glycolysis modulation were also confirmed by flow cytometry analysis (**Figure S2H-I**). In terms of the formation of MGCs (Orenstein, 2000; Verollet et al., 2015; Verollet et al., 2010), their numbers decreased or augmented under diminished (**Figure 3D and S2G**) or enhanced (**Figure 3D**) glycolysis, respectively. In line with these results, we observed at least a 50% decrease in HIV-1 infection and MGC formation in monocytes differentiated with TB-PE under diminished glycolysis condition (**Figure 3E-G**). To complement our pharmacological approach, we modified glycolysis by depriving the extracellular medium of glucose (**Figure 3H-K**). Accordingly, the lactate release by cmMTB-differentiated macrophages was reduced compared to the normal conditions (25 mM glucose) (**Figure 3I**). This deprivation had also a strong inhibitory effect on HIV-1 infection and MGC formation, without affecting cell density, in comparison to control cells (**Figure 3H and 3J-K**). Collectively, these results demonstrate that a shift towards aerobic glycolysis in our immunomodulatory macrophage model enhances its susceptibility to HIV-1 infection and the formation of infected MGCs.

**Figure 3.**
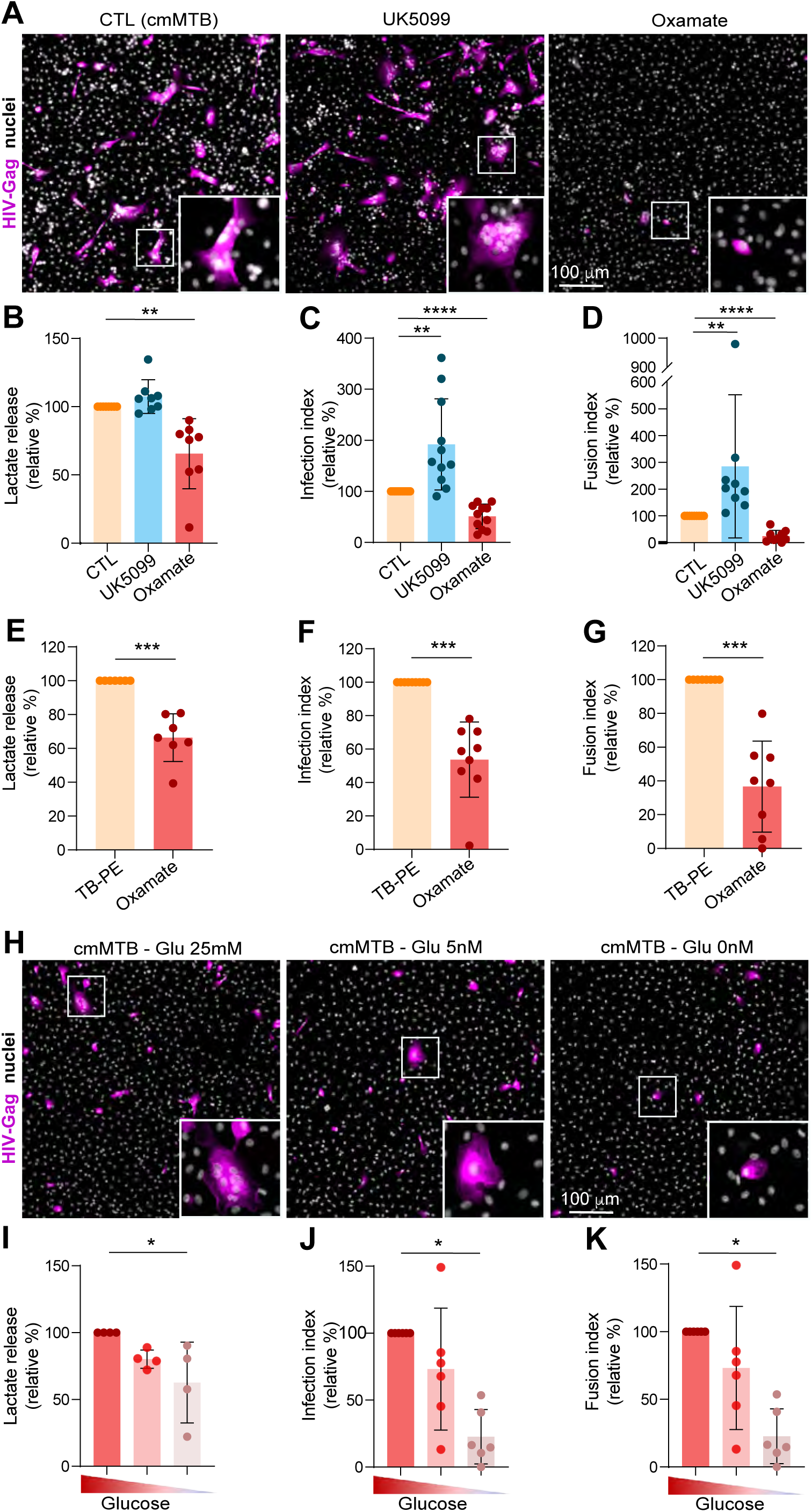
Modulation of aerobic glycolysis impacts HIV-1 infection of immunoregulatory macrophages. Monocytes from healthy subjects were differentiated in conditioned medium from Mtb-infected macrophages (cmMTB) or with pleural effusions (PE) from TB (TB-PE) for three days. At day two of differentiation, metabolic inhibitors were added to the culture medium. At day three, cells were infected by HIV-1 (NLAD8-VSVG) (for three further days) and HIV-1 infection of monocyte-derived macrophages (MDM) was measured (see experimental design, Supplemental Figure 2A-B). (**A-D**) Analysis of MDM infection upon UK5099 or oxamate treatment. (A) Representative IF images: HIV-Gag (magenta) and nuclei (DAPI, grey). Scale bar, 100 µm. (B) Lactate release measured at day three (24 h after drugs treatment); quantification of MDM infection index (C, n=9-11 donors) and MDM fusion index (D, n= nine donors), normalized to the control condition (CTL= cmMTB w/o treatment). (**E-G**) Analysis of HIV-1 infection of TB-PE-treated macrophages upon oxamate treatment by microscopy. (E) Lactate release measured at day three (24 h upon treatment); quantification of MDM infection index (F, n= nine donors) and MDM fusion index (G, n= eight donors), normalized to the control condition (CTL=cmMTB without treatment). (**H-K**) Analysis of MDM infection upon glucose deprivation (normal condition 25 mM, five mM or zero mM glucose) at day two. (H) Representative IF images: HIV-Gag (magenta) and nuclei (DAPI, grey). Scale bar, 100 µm. (I) Lactate release measured at day three (24 h upon treatment); quantification of MDM infection index (J, n= six donors) and MDM fusion index (K, n= six donors), normalized to the control condition (CTL = cmMTB without treatment). Mean +/- SD is shown. Each circle represents a single donor. Statistical analysis: data with normal distribution; *, p≤0.05; **, p≤0.01; *** p≤0.001; ****, p≤0.0001.

### TB-induced aerobic glycolysis promotes cell-to-cell dissemination of HIV-1 *via* TNT formation

Previous work demonstrated that TB-associated microenvironments alter HIV-1 dissemination through TNTs, which facilitate the transfer of the virus between macrophages, trigger their cell fusion, and lead to the formation of highly virus-productive MGCs. No other steps of the HIV-1 viral cycle were affected in these cells (Dupont et al., 2022; Dupont et al., 2020; Lastrucci et al., 2015; Souriant et al., 2019). To determine whether glycolysis regulates the cell-to-cell transfer of HIV-1, a co-culture was performed between uninfected recipient (labelled with CellTracker^+^) and HIV-1-infected donor (Gag^+^) cmMTB-differentiated macrophages for 24 h (**Figure S3A**). This timepoint allows sufficiently donor macrophages to transfer the virus to recipient cells primarily through fusion and mainly in a TNT dependent manner (Souriant et al., 2019). Accordingly, among HIV-1-Gag^+^ cells, about 35% were MGCs double positive for CellTracker (**Figure 4A-B**). Of note, infection by newly produced viruses is unlikely in 24 h. More importantly, both cell-to-cell transfer and fusion between these macrophages were significantly reduced when donor cells were under diminished glycolysis treatment prior to HIV-1 infection (**Figure 4A-B, left**). To further confirm that only the cell-to-cell transfer of the virus is affected by glycolysis, glycolysis was diminished specifically at the time of co-culture, which represents three days after HIV-1 infection of donor cells (**Figure S3A**). Under this condition, the number of double-positive cells was also decreased (**Figure 4B, right),** implying that glycolysis controls this mode of viral transmission without affecting earlier steps of the viral cycle.

**Figure 4.**
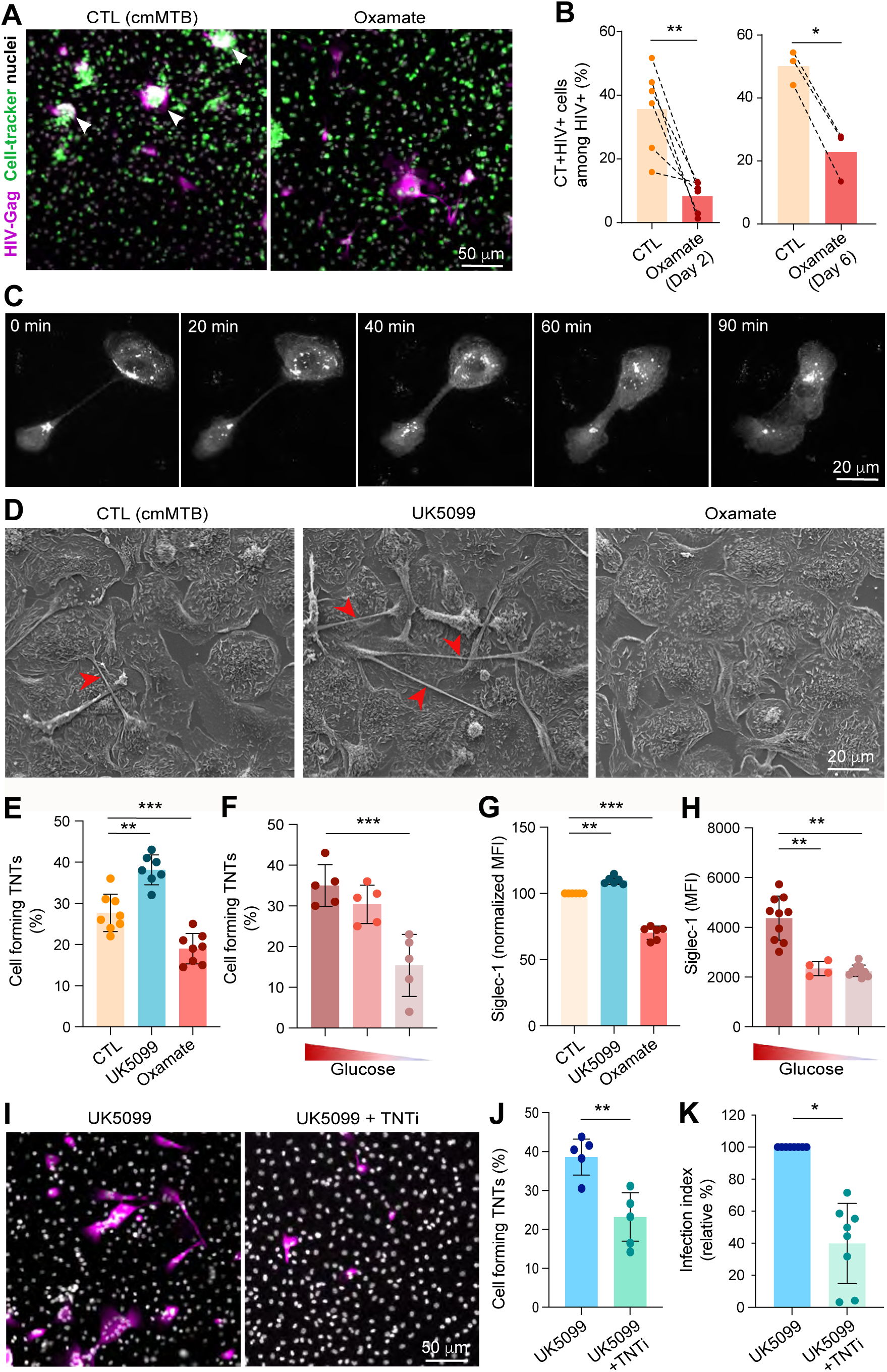
Glycolysis potentiates HIV-1 spread between macrophages. (**A-B**) Glycolysis favors HIV-1 transfer from one macrophage to another. (A) Representative IF images of cocultures (see experimental design, Supplemental Figure 3A). HIV-1 Gag (magenta), CellTracker (green) and DAPI (grey). Scale bar, 50 μm. Arrowheads show multinucleated double-positive cells (**B**) Quantification of the percentage of CellTracker^+^ cells among Gag^+^ cells in the condition of oxamate treatment at day two (left, n= six donors) or at day six (right, n= three donors). Histograms represent mean values. (**C**) Characterization of the fusion between two HIV-1 infected MDMs after tunneling-nanotube (TNT)-mediated connection. Conditioned medium from Mtb-infected macrophages (cmMTB)-treated cells were infected with a GFP-expressing viral strain (ADA-GFP-VSVG) at day three and live-microscopy was performed at day six (see also Movie 1). (**D-K**) Glycolysis regulates HIV-1-induced tunneling nanotube (TNT) formation and Siglec-1 surface expression, impacting HIV-1 dissemination in macrophages. CmMTB-treated cells were infected with HIV-1 at day three and TNT formation and Siglec-1 expression in MDMs were assessed at day six. (**D-F**) Analysis of TNT formation upon UK5099 or oxamate treatment in comparison with control condition (CTL =cmMTB without treatment). (D) Representative scanning electron microscopy (SEM) images. Scale bar, 20 µm. (E) Quantification of the percentage of cells forming TNTs (n= eight donors). (F) Analysis of TNTs formation after glucose deprivation (five mM or zero mM glucose) for two days in comparison with control condition (CTL=cmMTB without treatment; normal condition 25 mM glucose). Quantification of the percentage of cells forming TNTs (n= five donors, at least 200 cells/condition/donor). (**G-H**) Analysis of Siglec-1 expression at the surface of MDMs after UK5099/oxamate treatment (G) and glucose deprivation (H). Geomean fluorescence intensity (MFI) of Siglec-1 cell-surface expression, normalized to control condition (CTL = cmMTB without treatment) in D. (**I-K**) Analysis of HIV-1 infection of MDMs. CmMTB-treated cells were exposed to UK5099 and infected with HIV-1 in the presence or not of TNT inhibitor (TNTi, 20 μM). (I) Representative IF images: HIV-Gag (magenta) and nuclei (DAPI, grey). Scale bar, 50 µm. Quantification of TNT formation (J, n= five donors) and infection fusion index (K, n= eight donors), normalized to the control condition (cmMTB-treated cells in the presence of UK5099). Means +/- SD are shown. Each circle represents a single donor. Statistical analysis: data with normal distribution; *, p≤0.05; **, p≤0.01; ***, p≤0.001.

HIV-1 transfer between macrophages is usually mediated through TNTs, whereby HIV-1 can be found within the thicker, long and more stable versions of these structures, which are characterized by the presence of microtubules (Dupont et al., 2020). Using GFP-tagged virus and live imaging, thick TNTs containing HIV-1 were observed as a preliminary step to the fusion of infected macrophages (**Figure 4C and Movie 1**). To investigate the role of glycolysis in this process, we modulated its activity in HIV-1-infected cmMTB-differentiated macrophages based on our pharmacology approach. Scanning electron microscopy (SEM) analyses revealed that enhanced glycolysis significantly increased the number of cells forming TNTs, while diminished glycolysis led to a deficiency of cells forming these structures (**Figure 4D**). These observations were confirmed though an IF quantification approach (**Figure 4E and S3B**). Moreover, measurement of the number of thick and thin (lacking microtubules) TNTs (Dupont et al., 2020; Souriant et al., 2019) illustrated the importance of the glycolytic activity of macrophages for their formation; that is, enhanced glycolysis increased both types of TNT formation, while diminished glycolysis decrease them (**Figure S3C**). Importantly, TNT formation was also inhibited by reducing the level of glucose in the medium (**Figure 4F**). As Siglec-1, an I-type lectin receptor that recognizes and binds sialic acid-containing glycoproteins, stabilizes thick TNTs and enhances HIV-1 binding and transfer (Dupont et al., 2020), we investigated whether glycolysis affected its expression in our immunomodulatory macrophage model. Flow cytometry analyses revealed that the surface expression of Siglec-1 was reduced under diminished glycolysis or glucose deprivation, whereas it was increased under enhanced glycolysis (**Figure 4G-H**). Finally, as a proof-of-concept that TB-induced aerobic glycolysis exacerbates HIV-1 spread in macrophages through TNTs, these structures were pharmacologically hindered using a TNT inhibitor (TNTi) (Hashimoto et al., 2016; Souriant et al., 2019). Under enhanced glycolysis in cmMTB-differentiated macrophages, TNTi treatment significantly decreased both TNT formation and HIV-1 spread among cells (**Figure 4I-K**). Altogether, these findings demonstrate that TB-induced aerobic glycolysis in immunomodulatory macrophages enhances the formation of TNT, thus promoting the spread of HIV-1 among these cells.

## DISCUSSION

Macrophages play a pivotal role in HIV-1 dissemination and the establishment of persistent viral reservoirs across various host tissues, including the lungs, where they are the primary target for Mtb (Ganor et al., 2019; Hendricks et al., 2021; Rodrigues et al., 2017; Sattentau and Stevenson, 2016). The detrimental synergy between Mtb and HIV-1 significantly impacts the host, necessitating a deeper understanding of how TB-associated microenvironments enhance HIV-1 infection dynamics (Esmail et al., 2018). Our recent research has identified a specific immunomodulatory phenotype of macrophages, which is induced by TB-associated microenvironments and becomes highly susceptible to both Mtb and HIV-1 infections and replication. These macrophages are notably abundant in co-infection sites, such as the pulmonary pleural cavity in patients and the lung parenchyma in NHPs (Dupont et al., 2022; Dupont et al., 2020; Lastrucci et al., 2015; Souriant et al., 2019). The present study sheds light on how TB exacerbates HIV-1 infection in our immunomodulatory macrophage model, emphasizing the critical role of macrophage immunometabolism in the context of HIV-1/Mtb co-infection and how glycolysis is essential in TNT formation. Understanding this mechanism is crucial, as it as it highlights potential dangers for HIV patients receiving the TB vaccine and underscores the need to identify new targets for vaccines or therapies aimed at preventing TB-induced deleterious immunomodulatory macrophages.

Mtb is known to alter macrophage metabolism, but findings across studies vary due to differences in Mtb strains, timing of metabolic analysis, and macrophage origins (Cumming et al., 2018; Kumar et al., 2019; Llibre et al., 2021; Olson et al., 2021). In general, it is well-accepted that infected macrophages undergo aerobic glycolysis as part of the Warburg effect to eliminate Mtb and other intracellular pathogens (Lugo-Villarino and Neyrolles, 2014). This shift towards glycolysis is evident in studies showing GlycoATP production boosts in macrophages challenged with non-infectious Mtb strains (Gleeson et al., 2016; Marin Franco et al., 2020). Many studies report increased glycolysis post-infection with virulent Mtb strains, but often do not distinguish between infected and bystander cells (Braverman et al., 2016; Huang et al., 2019; Marin Franco et al., 2020; Russell et al., 2019; Shi et al., 2019). The use of TB-associated microenvironments enables the study of the bystander effect induced by infected cells on the metabolic state of their neighbor cells, including recruited circulating monocytes at the infection site. Using *in vitro* (cmMTB) and *ex vivo* (PE-TB) models to mimic TB-associated microenvironments, we observed increased glycolysis in monocytes differentiating into immunomodulatory macrophages, as measured by metabolic analyses, an enriched glycolytic gene signature, and activation of the metabolic regulator HIF-1α. Although treating monocytes with CmMTB or TB-PE leads to an immunomodulatory macrophage phenotype dependent on the IL-10/STAT3 (Lastrucci et al., 2015; Souriant et al., 2019) and type-I interferon (IFN-I)/STAT1 (Dupont et al., 2022; Dupont et al., 2020) signaling pathways, the literature well-documents that IL-10 and IFN-I are associated with the suppression of glycolytic activity (Olson et al., 2021; Yeh et al., 2018). This evidence suggests that these cytokines alone are unlikely responsible for the metabolic changes observed in macrophages. We infer that mycobacterial antigens in the fluids might drive glycolysis in the cells, as attenuated Mtb strains consistently induce glycolysis in macrophages, unlike the live and virulent ones. Although this work did not focus on the impact of the glycolytic shift on Mtb control, it may contribute to high intracellular replication of the bacillus reported in these permissive macrophages (Lastrucci et al., 2015). This supports the notion that increased glycolysis does not necessarily improve infection control. Recent studies have shown that monocytes from TB patients, inherently biased toward glycolysis, differentiate into dendritic cells that do not engage in glycolytic flux effectively, leading to poor migratory capacities (Maio et al., 2024). Thus, despite this shift towards increased glycolysis, prior exposure to inflammatory signals may render macrophages unable to engage the strong and lasting glycolytic reprogramming seen in inflammatory M1 macrophages controlling intracellular Mtb replication. Future investigations shall address this important issue and establish *in vivo* correlations between the glycolytic shift and the macrophage compartment in biopsies and samples from NHP and TB patients. However, modulation of glycolytic activity in alveolar and interstitial macrophages in Mtb-infected mice has already shown importance in controlling bacterial growth (Huang et al., 2019).

Viruses lack the metabolic machinery for their survival and have developed strategies to exploit the metabolic resources of their host cells, as reported for HIV-1 (Saez-Cirion and Sereti, 2021). Our knowledge of these metabolic adaptations mainly comes from the study of CD4^+^ T lymphocytes, which are the most researched host cells for this virus. HIV-1 fitness is favored in T lymphocytes with high metabolic activity (Shehata et al., 2017; Valle-Casuso et al., 2019). The metabolic program of different T cell subtypes is crucial in determining their susceptibility to infection, regardless of their activation level, with glycolysis and glutaminolysis being key to sustaining the pre-integration steps of HIV-1 infection (Clerc et al., 2019; Loisel-Meyer et al., 2012; Palmer et al., 2016; Shehata et al., 2017; Valle-Casuso et al., 2019). Under specific conditions of TB-associated microenvironments, we observed that early steps of the viral cycle are not affected by the inhibition of glycolysis in macrophages. However, the metabolic activity of CD4^+^ T cells is critical not only in facilitating intracellular replication during HIV-1 infection but also in governing the overall infection process (Shehata et al., 2017). For myeloid cells, previous studies have shown that CD16^+^ monocytes, the most permissive monocyte subset to HIV-1 infection (Ellery et al., 2007; Rodrigues et al., 2017), exhibit heightened expression of Glut-1, increased glucose uptake, and elevated lactate release (Palmer et al., 2016). However, a direct link between glycolysis and the susceptibility of monocyte/macrophages to HIV-1 was missing.

In this study, using drugs to selectively modulate the glycolytic pathway, we demonstrated that this pathway is involved in HIV-1 infection and the formation of MGCs, hallmarks of HIV-1 infection of macrophages (Han et al., 2022; Verollet et al., 2015). In TB-associated microenvironments, oxamate, a competitive inhibitor of Lactate Dehydrogenase A (LDHA), reduced the number of infected MGCs by more than two-fold. Oxamate was preferred over 2-deoxyglucose (2-DG) due to the latter’s controversial specificity and relevance as a glycolysis inhibitor (Saez-Cirion and Sereti, 2021). Oxamate also blocks lactate production without altering pyruvate’s role as a fuel for mitochondrial respiration, distinguishing the contribution of glucose to ATP production by glycolysis *versus* OXPHOS. Complementing the use of pharmacological inhibitors, glucose deprivation experiments confirmed our results. Similar findings have been observed in previous studies, highlighting the importance of macrophage glycolytic activity in viral replication for viruses like dengue (Fontaine et al., 2015) and murine norovirus (Passalacqua et al., 2019). Interestingly, within the context of vesicular stomatitis virus (VSV) infection, glycolysis promotes viral replication by negatively regulating IFN-I and antiviral responses (Zhang et al., 2019). Since immunoregulatory macrophages exhibit a strong but defective IFN-I signature (Dupont et al., 2022), it is plausible that glycolysis may impair the IFN-I anti-viral, and *vice-versa* (Maio et al., 2024). To explore whether environmental triggers of glycolysis influence HIV-1 infection in macrophages and broader HIV-1 pathogenesis, additional research is needed beyond the context of Mtb co-infection. Current knowledge indicates that HIV-1 infection can enhance glucose metabolism in macrophages (Datta et al., 2016). However, this effect may vary depending on the macrophage type, state, and infection timing (Castellano et al., 2019; Hollenbaugh et al., 2011). Although we did not examine the metabolic status of macrophages post-HIV-1 infection alone, sustained glycolytic activity might optimize long-term HIV-1 infection. Another aspect to consider is how macrophage polarization impacts susceptibility to HIV-1 (Cassol et al., 2010; Cassol et al., 2009; Hendricks et al., 2021). Interestingly, extreme M1 and M2 polarized macrophages, which have distinct metabolic profiles, are less susceptible to HIV-1 infection compared to unpolarized macrophages (Saha et al., 2017). However, these studies primarily used cell-free viral infection, which is less common *in vivo* than cell-to-cell infection (Bracq et al., 2018; Han et al., 2022; Mascarau et al., 2023). Future research should focus on the link between metabolism and heterotypic intercellular HIV-1 infection, especially the transfer from T cells to macrophages, which is influenced by macrophage polarization (Mascarau et al., 2023).

As aforementioned, the cell-to-cell transfer of HIV-1 is more efficient than infection with cell-free viruses and plays a critical role in virus dissemination *in vivo*, especially in macrophages (Dupont and Sattentau, 2020; Han et al., 2022; Murooka et al., 2012; Sewald et al., 2012). The primary mechanism for this transfer involves TNTs, which facilitate the transport of viral particles. TNT formation is exacerbated in TB-associated microenvironments enriched with IL-10. However, the molecular mechanisms behind TNT formation are not fully understood (Dupont et al., 2020; Eugenin et al., 2009; Hashimoto et al., 2016; Lotfi et al., 2020; Okafo et al., 2017; Souriant et al., 2019; Uhl et al., 2019). We showed that TNT formation between infected macrophages not only contributes to their fusion into MGCs but also that glycolysis plays a crucial role in controlling the formation of these structures. Blocking glycolysis reduces TNT formation, while promoting this metabolic pathway increases their presence. TNTs are F-actin-based open-ended membranous channels connecting cells over various distances. Their formation and stability can be influenced by extracellular conditions, such as nutritional deprivation, oxidants, acidic conditions, and cytokines (Goodman et al., 2019; Lou et al., 2012). Our work highlights the role of intracellular metabolism, particularly glycolysis, in promoting TNT formation in macrophages. However, whether these findings can be translated to other cell types that form TNTs remains an open question, as studies in cancer cells and mesenchymal stem cells have shown contradictory results in TNT formation and dynamics (Liu et al., 2014; Thayanithy et al., 2014).

In macrophages, two types of TNTs exist: thin TNTs (<0.7 μm in diameter, containing F-actin) and thick TNTs (>0.7 μm in diameter, rich in F-actin and microtubules) (Onfelt et al., 2006; Souriant et al., 2019). Glycolysis modulation affects both types, with increased glycolytic activity in immunoregulatory macrophages potentially providing the energy needed for actin cytoskeletal rearrangements essential for TNT formation. ATP is crucial for supporting cellular functions involving actin remodeling, such as cell migration and epithelial to mesenchymal transition (DeWane et al., 2021). In HIV-1 infected macrophages, ATP is also vital for the release of particles from virus-containing compartments (Graziano et al., 2015). Our study suggests that glycolysis, by promoting TNT formation, enhances HIV-1 dissemination between macrophages. Besides HIV-1, thick TNTs can transfer various organelles, including mitochondria, which can alter the metabolism and functional properties of recipient cells (Goliwas et al., 2023; Hekmatshoar et al., 2018; Wang et al., 2021), adding complexity to the model. Within the context of thick TNTs, Siglec-1 (a sialic acid-binding lectin) plays a critical role in their stabilization (Dupont et al., 2020). We show that the expression of Siglec-1 is influenced by glycolytic activity within macrophages. High levels of glycolysis upregulate Siglec-1 expression, enhancing the formation and stability of thick TNTs, thereby promoting efficient viral dissemination. Given that Siglec-1 is an IFN-stimulated gene (Dupont et al., 2022), it is plausible that glycolysis influence that capacity of bystander macrophages to produce autocrine IFNβ (Olson et al., 2021). This metabolic regulation underscores the significance of glycolysis in modulating macrophage functions and intercellular communications, particularly in the context of viral infections. A deeper understanding of cell-to-cell communication mechanisms mediated by TNTs and the influence of metabolism on these processes is still needed.

To conclude, our study demonstrates that TB-associated microenvironments induce aerobic glycolysis in immunoregulatory macrophages, increasing their propensity to form TNTs and facilitating HIV-1 transfer between macrophages. The role of glycolysis during viral infection remains unclear, especially regarding its potential dysregulation in the context of co-infection with other pathogens. Recent reports indicate that glycolysis in tissue macrophages, which exhibit an intermediate M1/M2 profile, is necessary for HIV-1 reactivation from latency (Real et al., 2022). This suggests glycolysis could be a potential target in future HIV-1 eradication strategies. Our findings support the idea that targeting glycolysis could disrupt viral progression in the context of Mtb co-infection. Further research is necessary to fully comprehend the metabolic effects mediated by TB-associated microenvironments on the lung macrophage compartment. We speculate that the identification of the mycobacterial antigen in the TB-associated milieu driving glycolysis would allow an extremely targeted therapy to specifically inhibit this metabolic rewiring in human macrophages. This will provide opportunities to disentangle the complex beneficial and detrimental roles that the glycolysis/OXPHOS balance plays in immunity and co-infection with HIV-1.

## MATERIALS AND METHODS

### Human subjects

Human primary monocytes were isolated from healthy subject (HS) buffy coat (provided by Etablissement Français du Sang, Toulouse, France, under contract 21/PLER/TOU/IPBS01/20,130,042) and differentiated towards macrophages (Souriant et al., 2019). According to articles L12434 and R124361 of the French Public Health Code, the contract was approved by the French Ministry of Science and Technology (agreement number AC 2009921). Written informed consents were obtained from the donors before sample collection.

PE samples from patients with TB were obtained by therapeutic thoracentesis by physicians at the Hospital F. J Muñiz (Buenos Aires, Argentina). The diagnosis of TB pleurisy was based on a positive Ziehl–Nielsen stain or Lowestein–Jensen culture from Pe and/or histopathology of pleural biopsy and was further confirmed by an Mtb-induced IFN-γ response and an adenosine deaminase-positive test (Light, 2010). Exclusion criteria included a positive HIV test, and the presence of concurrent infectious diseases or non-infectious conditions (cancer, diabetes, or steroid therapy). None of the patients had multidrug-resistant TB. The research was carried out in accordance with the Declaration of Helsinki (2013) of the World Medical Association and was approved by the Ethic Committee of the Hospital F. J Muñiz (protocol number: NIN-2601-19). Written informed consent was obtained before sample collection.

### Bacteria

Mtb (H37Rv, see supplemental information, Table 1) was grown at 37°C in Middlebrook 7H9 medium, supplemented with 10% albumin-dextrose-catalase, as described previously (Lastrucci et al., 2015). Exponentially growing Mtb was centrifuged (460g) and resuspended in PBS (MgCl_2_, CaCl_2_ free, Gibco). Clumps were dissociated by 20 passages through a 26-G needle and then resuspended in RPMI-1640 containing 10% FBS. Bacterial concentration was determined by measuring the optical density (OD) at 600 nm.

### Viruses

Viral stocks were generated by transient transfection of 293T cells with the proviral plasmids coding for HIV-1 NLAD8, kindly provided by Serge Bénichou (Institut Cochin, Paris, France), with VSV-G envelope, as previously described (Mascarau et al., 2023) (Table 1). Supernatants were harvested 48 h post-transfection and p24 antigen concentration was assessed by a homemade ELISA. HIV-1 infectious units were quantified using TZM-bl cells as previously reported (Mascarau et al., 2023).

### Human monocyte-derived macrophage culture

Monocytes were isolated and differentiated towards monocyte-derived macrophages as described (Lastrucci et al., 2015; Souriant et al., 2019). Briefly, peripheral blood mononuclear cells were recovered by gradient centrifugation on Ficoll-Paque Plus (GE Healthcare). CD14^+^ monocytes were then isolated by positive selection magnetic sorting, using human CD14 Microbeads and LS columns (Miltenyi Biotec). Cells were then plated at 1.6×10^6^ cells in 6-well plates and allowed to differentiate for five to seven days in RPMI-1640 medium (GIBCO), 10% Fetal Bovine Serum (FBS, Sigma-Aldrich) and human M-CSF (20 ng/mL, Peprotech) before infection with Mtb H37Rv for conditioned-media preparation. The cell medium was renewed every third or fourth day.

### Preparation of conditioned media of Mtb-infected macrophages

CmMTB) was prepared as reported previously (Lastrucci et al., 2015). Briefly, MDM were infected with Mtb H37Rv at a MOI of three. After 18 h of infection at 37°C, culture supernatants were collected, filtered by double filtration (0.2 µm pores) and aliquots were stored at −80°C. CmCTR was obtained from uninfected macrophages.

### Treatment of monocytes with the secretome of Mtb-infected macrophages or Pleural effusion from TB patients

Freshly isolated CD14^+^ monocytes from HS were allowed to adhere in the absence of serum (4×10^5^ cells in 50 μL in 24-wells plates or 2×10^6^ cells in 1.5 mL in 6-wells plates). After one h, cmMTB or cmCTR supplemented with 20 ng/mL M-CSF and 20 % FBS were added to the cells (vol/vol). For experiments with Pe, samples were collected in heparin tubes and centrifuged at 300 g for ten min at room temperature. The cell-free supernatants were transferred into new plastic tubes, further centrifuged at 12000 g for ten min and aliquots were stored at −80°C. After having the diagnosis of the PE samples, pools were prepared by mixing same amounts of individual PE associated to a specific etiology. The pools were de-complemented at 56°C for 30 min and filtered through 0.22 μm pores to remove any remaining debris or residual bacteria. A pool of PE samples from ten patients with active TB was prepared. Additionally, a pool of PE from five patients with transudates secondary to heart failure was included as control. Both pools were supplemented with 40 ng/mL M-CSF and 40% FBS were added to the cells (25 % vol/vol). Cells were then cultured for three days. Cell-surface expression of macrophage activation markers was measured by flow cytometry using standard procedures.

### HIV-1 infection

At day three of differentiation, 0.5×10^6^ macrophages in 24-wells plates were infected with HIV-1 VSVG (or HIV-Gag-iGFP-VSVG for live imaging) strain at MOI of one in a fresh culture medium. HIV-1 infection and replication were assessed at day three post-infection by measuring p24-positive cells by immunostaining or flow cytometry, as described previously (Mascarau et al., 2020).

### Co-culture assays

Half of the cmMTB-treated macrophages was infected with HIV-1 NLAD8 VSVG for three days. At day six, the other half of the cell population was stained with CellTracker Green CMFDA Dye (Thermo Fisher Scientific). Thereafter, cells were washed three times with PBS Mg2^+^/Ca2^+^ and detached using trypsin 0.25% EDTA (Gibco) for 15 min. Then, they were co-cultured at a 1:1 ratio on glass coverslips in 24-well for 24 h. They were fixed with PFA 3.7 %, sucrose 15 mM in PBS during one h, and HIV-1 transfer was assessed by immunofluorescence as described previously (Souriant et al., 2019). In defined experimental kinetic conditions, oxamate was added 24 h before HIV-1 infection (day two) or 30 min after coculture (day six).

### Drug treatments and glucose deprivation experiments

Two days after purification, monocytes were treated with 20 mM sodium oxamate (Santa Cruz Biotechnology), 50 μM UK5099 (MedChem Express) or 60 μM GSK 2837808A (MedChem Express). For the experiments using TNTi (Pharmeks), cells were treated with 20 μM TNTi at days zero and three, as described previously (Souriant et al., 2019). For glucose deprivation experiments, the medium was changed at day two of differentiation with RPMI 1640 Medium without glucose (Fisher Scientific) and FBS complemented with 0.5 or 25 mM of glucose (Thermo Scientific).

### Immunofluorescence microscopy

Cells were fixed with PFA 3.7% and sucrose 15 mM in PBS. Cells were permeabilized with Triton X-100 0,3% for ten min, and saturated with PBS BSA one % for 30 min. See Table 1 for antibodies and reagents. Cells were incubated with anti-HIV-1-Gag KC57 antibody RD1 (1:100) in PBS BSA one % for one h, washed and then incubated with Alexa Fluor 555 Goat antiMouse IgG secondary antibody (1:1000), Alexa Fluor 488 Phalloidin (1:500) or WGA 488 and DAPI (500 ng/mL) in PBS BSA one % for 30 min. For microtubule staining, cells are incubated with anti-tubulin antibodies (1:100, Sigma). For HIF1-α staining, anti-HIF1-α antibody (1:50) was used. Coverslips were mounted on a glass slide using Fluorescence Mounting Medium (Dako). Images for quantification of infection were acquired using a Zeiss Axio Imager M2 and a 20×/0.8 Plan Apochromat or 40×/0.95 Plan Apochromat objectives (Zeiss). Images were acquired and processed using the Zeiss Zen software ORCA-flash 4.0 LT (Hamamatsu) camera. For confocal images (analysis of TNTs), specimens were observed with a Zeiss LSM710 confocal microscope that uses a Zeiss AXIO Observer Z1 inverted microscope stand with transmitted (HAL), UV (HBO), and laser illumination sources. Images were acquired with a Zeiss ×63 (oil) NA 1.35 objective. For all images, visualization and analysis was performed with ImageJ. The HIV infection index (total number of nuclei in HIV-stained cells divided by total number of nuclei × 100) was quantified, as described in (Souriant et al., 2019). The fusion index is defined as the number of nuclei present in a multinucleated giant cell (>two nuclei) relative to the total number of nuclei (Verollet et al., 2015). For TNT quantification, TNTs were detected and counted using F-actin and microtubule staining. Thick and thin nanotubes were quantified: thin membrane nanotubes contained only F-actin, whereas thick TNTs contained both F-actin and microtubules, as described in (Souriant et al., 2019).

### Flow cytometry

Adherent cells were harvested after 15 min of incubation in trypsin 0.05 % EDTA (Gibco) and washed with PBS (Gibco). After five min centrifugation at 1300 rpm, pellets were resuspended in cold staining buffer (PBS, 2 mM EDTA, 0.5% FBS) with fluorophore-conjugated antibodies (Table 1). For intracellular staining, cells were fixed with PFA 3.7% for one h and stained with fluorophore-conjugated antibodies (Table 1) in staining buffer with 0.15% triton. After staining, cells were washed with cold staining buffer, centrifuged for five min at 1400 rpm at 4°C, and analyzed by flow cytometry using BD LSRFortessa flow cytometer (BD Biosciences, TRI Genotoul plateform) and the associated BD FACSDiva software. Data were then analyzed using the FlowJo_V10 software (FlowJo, LLC). For mitochondria analysis, adherent cells were stained for ten min at 37°C with Mitotracker or MitoSOX, according to supplier protocols. For glucose uptake, cells were incubated with the fluorescent glucose analog 2-(N-(7-Nitrobenz-2-oxa-1,3-diazol-4-yl)-Amino)-2-Deoxyglucose (2-NBDG) (10 μM, Invitrogen, California, USA) in PBS for 30 min. Thereafter, cells were washed and intracellular 2-NBDG was measured by flow cytometry.

### Determination of lactate release

Lactate production concentrations in culture media were measured using the spectrophotometric assays Lactate Kit from Wiener (Argentina), which are based on the oxidation of lactate (Marin Franco et al., 2020). The absorbance was read using CLARIOstar Microplate Reader and its software.

### Transmission electron microscopy

Cells were fixed in 2.5 % glutaraldehyde and two % paraformaldehyde (EMS, Delta-Microscopies) dissolved in 0.1 M Sorensen buffer (pH 7.2) during two h at room temperature, and then preserved in one % PFA dissolved in Sorensen buffer. Adherent cells were treated for one h with one % aqueous uranyl acetate then dehydrated in a graded ethanol series and embedded in Epon. Sections were cut on a Leica Ultracut microtome and ultrathin sections were mounted on 200 mesh onto Formvar carbon-coated copper grids. Finally, thin sections were stained with one % uranyl acetate and lead citrate and examined with a transmission electron microscope (Jeol JEM-1400) at 80 kV. Visualization and quantification of mitochondria was performed as described previously (Genoula et al., 2020). Images were acquired using a digital camera (Gatan Orius).

### Scanning electron microscopy

Cells were washed three times for five min in 0.2 M cacodylate buffer (pH 7.4), post-fixed for one h in 1% (wt/vol) osmium tetroxide in 0.2 M cacodylate buffer (pH 7.4) and washed with distilled water. Samples were dehydrated through a graded series (25–100%) of ethanol, transferred in acetone, and subjected to critical point drying with CO_2_ in a Leica EM CPD300. Dried specimens were sputter-coated with three nm platinum with a Leica EM MED020 evaporator and were examined and photographed with a FEI Quanta FEG250.

### Real-Time cell metabolic analysis using SeaHorse®

Cells (5×10^5^ cells/well) were plated after isolation on XFe-24-cell culture plates (Agilent), treated with cmCTR/cmMTB/HF-PE or TB-PE for one, two or three days. One h before the assay, images of each well were taken (Incucyte) to allow normalization based on the area of the well occupied by the cells. Then, cells were washed and replaced with Dulbecco’s Modified Eagle’s Medium (DMEM) (Sigma-Aldrich) supplemented with 4.5 g/L d-glucose, two mM glutamine and two mM pyruvate, following by an incubation without CO_2_ at 37 °C for 40 min. Mito Stress assay was performed by sequential addition of 1.5 μg/mL oligomycin (inhibitor of ATP synthesis), 0.7 μM carbonyl cyanide 4-(trifluoromethoxy) phenylhydrazone (FCCP, uncoupling agent) and one μM rotenone/antimycin A (inhibitors of complex I and complex III of the respiratory chain, respectively). ATP rate test was performed by sequential addition of 1.5 μg/mL oligomycin (inhibitor of ATP synthesis) and one μM rotenone/antimycin A (inhibitors of complex I and complex III of the respiratory chain, respectively).

### Live imaging

For live imaging, three days after HIV-1-GFP infection, specimens were observed on an Andor/Olympus spinning disk microscope equipped with a Yokogawa CSU-X1 scanner unit and an emCCD camera (Andor iXon 888) under the control of the iQ3 software (Andor Oxford Instruments company). Images were acquired with an Olympus ×60 (oil) NA 1.35 objective for DIC and GFP (HIV) signal. One image every minute and a half.

### Transcriptomics and GSEA analysis

The transcriptomic data of cells conditioned with cmCTR and cmMTB supernatants for three days was described in (Dupont et al., 2022) and available as GEO accession number: GSE139511. We pre-processed the data as in the original publication and applied GeneSet Enrichment Analysis (GSEA) using the Hallmark genesets available in MSigDB (v7.5.1) in order to gain insights in the pathways differentially regulated between the two conditions (Subramanian et al., 2005). GSEA allows to statistically test whether a geneset is significantly enriched when comparing two conditions based on their expression profiles. We used the following parameters: Metric for ranking genes: Signal2Noise, Permutation type: gene_set, Number of permutations: 1000.

### Analysis of extracellular metabolites by proton ^1^H-NMR

Culture supernatant (180 µL) of CD14+ monocytes was collected at different time points (12, 20, 40 and 60 h) after treatment with the secretome of Mtb-infected macrophages. 20 µL of ten mM (trimethylsilyl)propionic acid d4 (TSPd4) solution dissolved into D2O were added to samples for frequency calibration and concentration measurements. The final volume of 200 µL of the resulting samples were transferred into three mm NMR tubes. Samples were analyzed by 1H-1D NMR on a Bruker Avance III HD 800 MHz spectrometer equipped with a five-mm quadruple-resonance QCI-P (H/P-C/N/D) cryogenically cooled probe head. NMR spectra were recorded and processed using Bruker TopSpin 3.2. 1H-1D NMR spectra were acquired using a quantitative zgpr30 sequence at 280 K with 32 scans, 131k points, an acquisition time of four sec, and an recycle delay of eight sec. Lactate and glucose consumption fluxes are shown.

### Statistical analysis

All statistical analyses were performed using GraphPad Prism 9 (GraphPad Software Inc.). Two-tailed paired or unpaired t-test was applied on data sets with a normal distribution (determined using Kolmogorov-Smirnov test), whereas two-tailed Mann-Whitney (unpaired test) or Wilcoxon matched-paired signed rank tests were used otherwise. Bar histograms represent mean with standard deviation (SD) for data with a normal distribution, and median with interquartile range otherwise. When multiple comparisons were done, the statistical analyses used were detailed in the corresponding Figure legend. p<0.05 was considered as the level of statistical significance (*, p≤0.05; **, p≤0.01; *** p≤0.001; **** p≤0.0001).

## ACKNOWLEDGEMENTS

We greatly acknowledge Flavie Moreau and Céline Berrone, IPBS and Genotoul Anexplo-IPBS, for accessing the BSL3 facilities; and the Genotoul TRI-IPBS facilities for imaging and flow cytometry, in particular Emmanuelle Näser, Serge Mazères and Eve Pitot. We also greatly acknowledge Serge Bénichou for critical reading of the manuscript. This work was supported by the *Centre National de la Recherche Scientifique*, *Université Paul Sabatier*, the *Institut National de la Santé et de la Recherche Médicale*, the *Agence Nationale de la Recherche* (ANR16-CE13-0005-01, ANR DFG 2020 JA-3038/2-1), the *Agence Nationale de Recherche sur le Sida et les hépatites virales (ANRS I MIE)* grant numbers: ECTZ 118551/118554, ECTZ 205320/305352, ECTZ 242543/293306, *Sidaction*, and the *Fondation pour la Recherche Médicale*. Part of this work was also supported by grants from the ANRS (ECTZ 88611 and 192763) to FB. QH is a recipient of a UM-CBS2 doctoral fellowship. We also thank the AIDS Research and Reference Reagent Program, Division of AIDS, NIAID. S.M and N.F are PhD candidates at Université Toulouse III Paul Sabatier supported by ANRS. Z.V was supported by ANR.

---

**Supplemental Figure 1.**
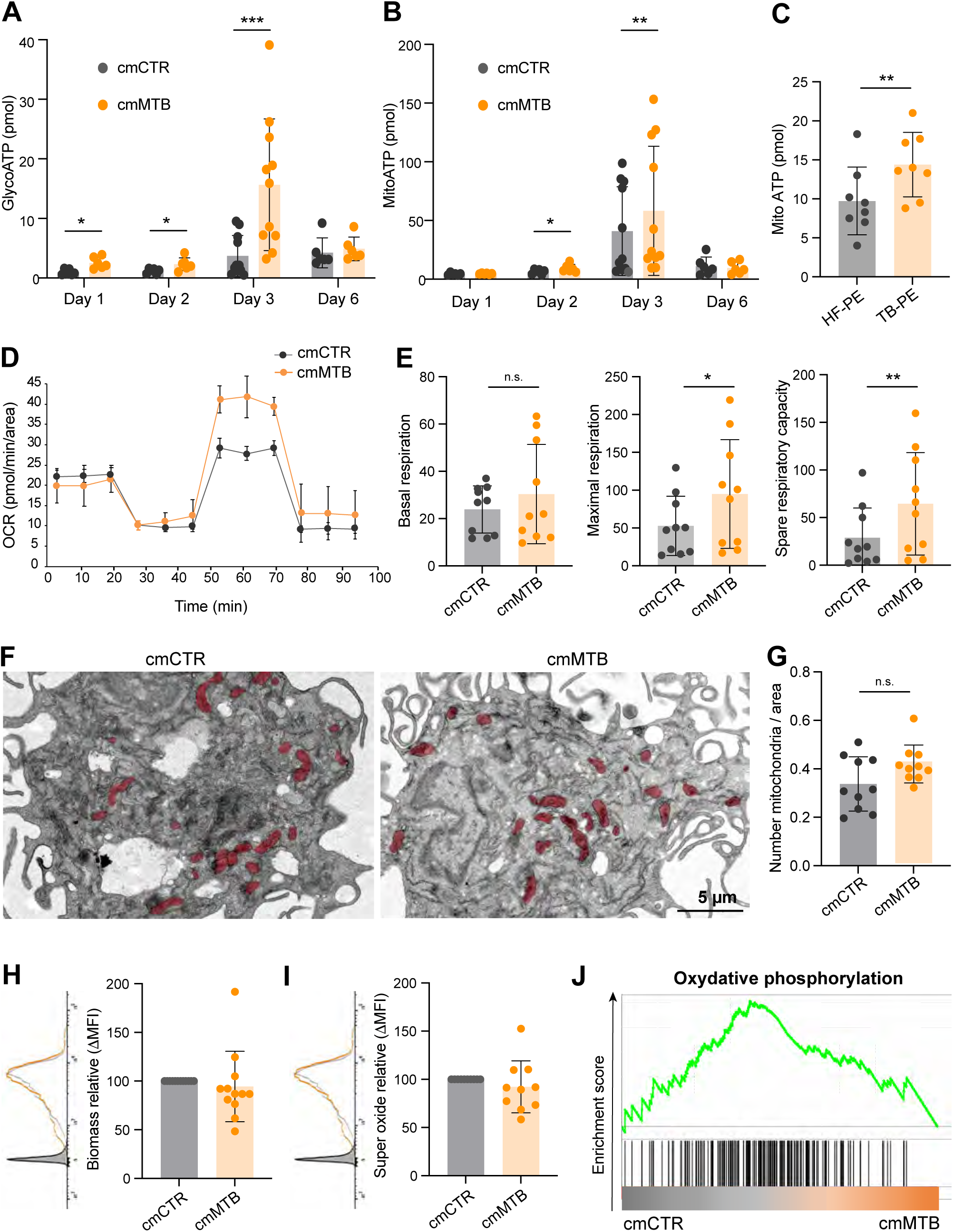

**Supplemental Figure 2.**
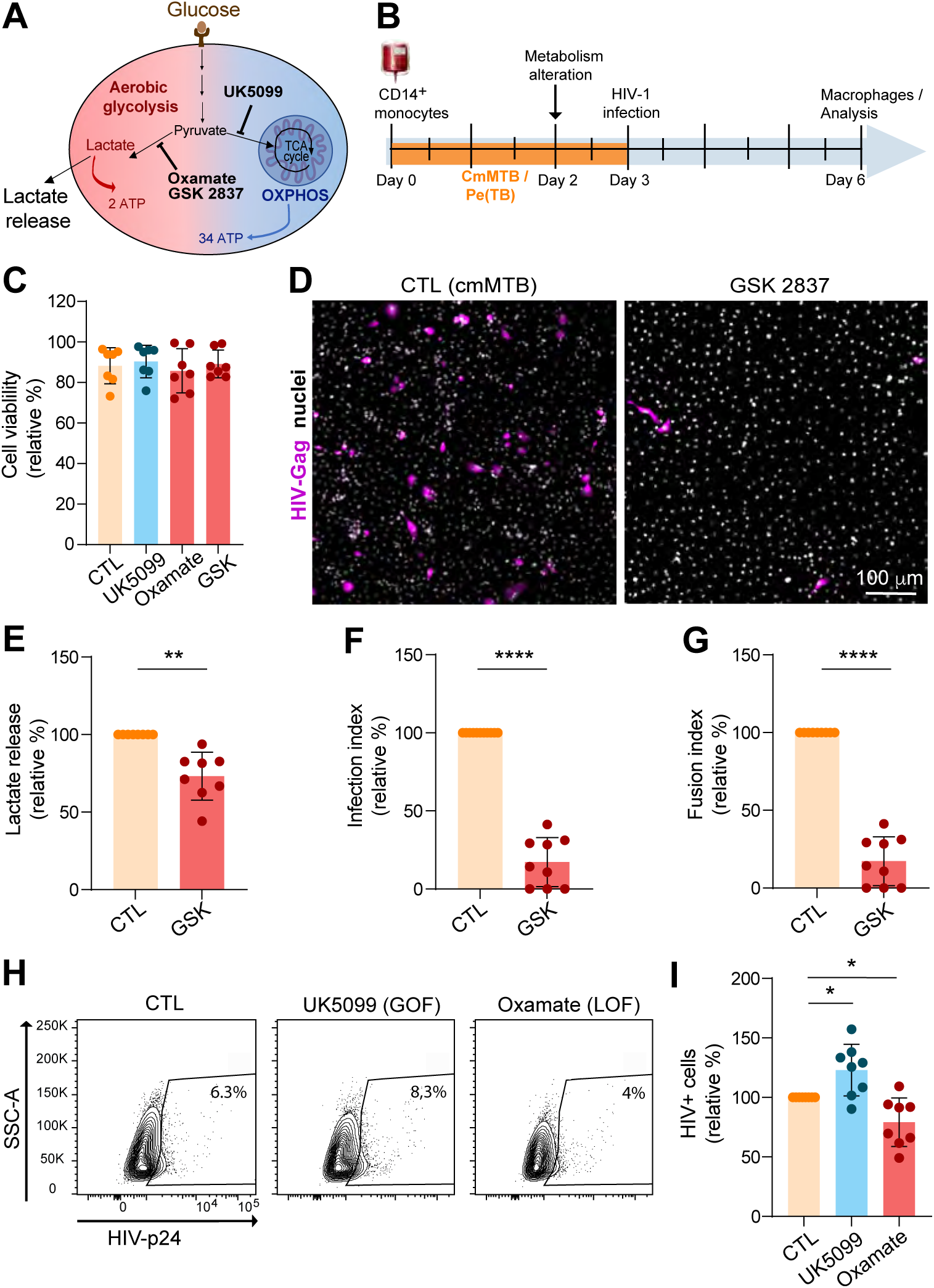

**Supplemental Figure 3.**
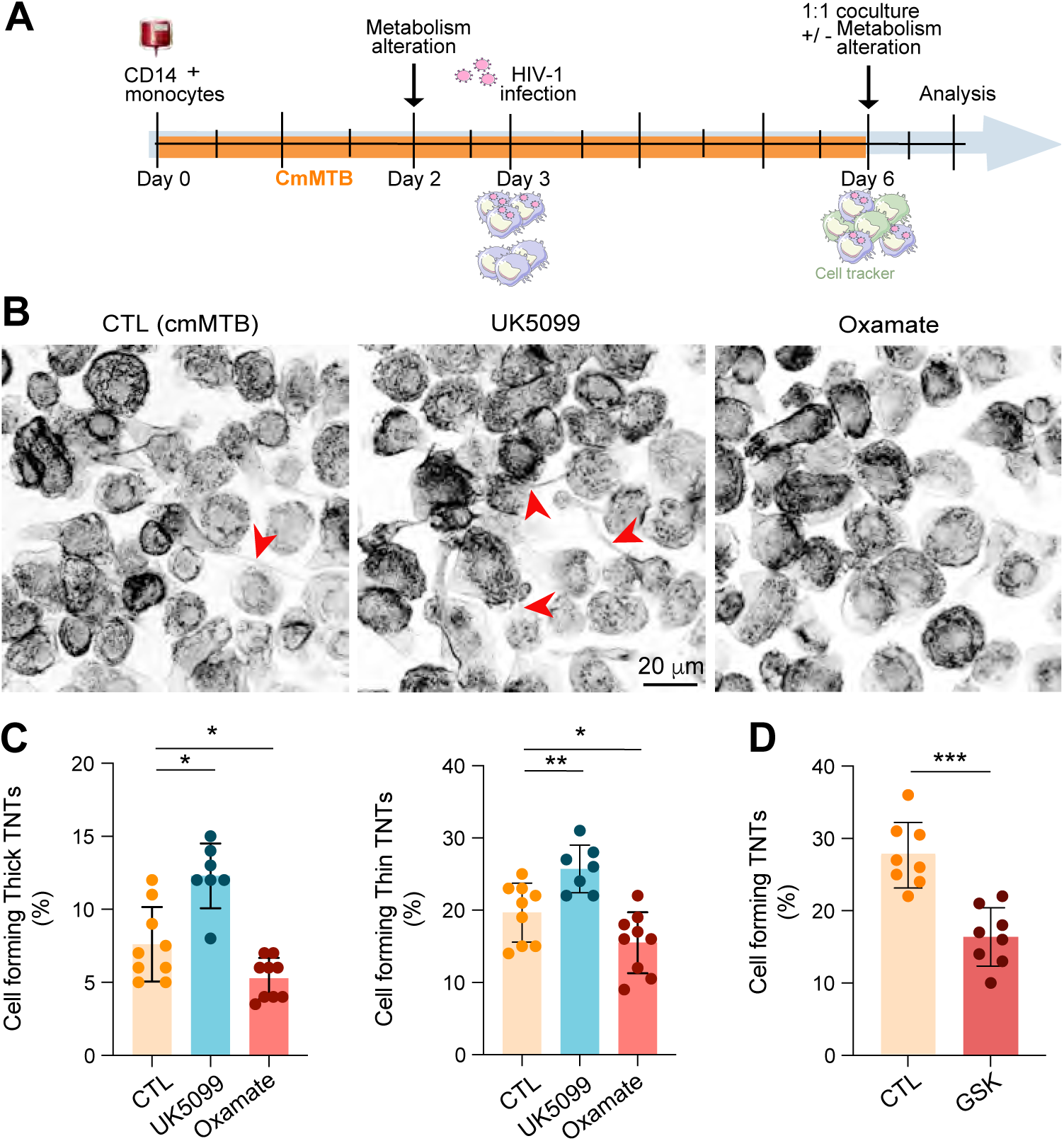

