## Supplemental Information Text for "Inhibition of glycolysis in tuberculosis-mediated metabolic rewiring reduces HIV-1 spread across macrophages"

### Vahlas et al, Supplemental Information

### Supplemental Figure legends

**Supplemental Figure 1** (related to Figure 1)**. Metabolic analysis of macrophages in TB-derived environments.**

Monocytes from healthy subjects were treated either with conditioned medium from mock- (cmCTR, grey) or Mtb-infected macrophages (cmMTB, orange), or with pleural effusions (PE) from TB (TB-PE, orange) or heart failure (HF-PE, grey) patients. (**A-E**) Analysis of metabolic parameters using an Agilent Seahorse XFe24 Analyzer. (A-B) Kinetics of total GlycoATP (**A**) or MitoATP (**B**) production in cmCTR and cmMTB treated macrophages. (**C**) Total MitoATP production in TB-PE- and HF-PE-treated macrophages at day three. (**D**) Representative experiment of mitostress assay showing the evolution of the oxygen consumption rate (OCR) following sequential injection of drugs at day three. (E) Quantification of basal respiration (left), maximal respiration (middle) and spare respiratory capacity (right) (pmol/min/area) at day three. (**F-G**) Analysis of the number of mitochondria per cell area between cmCTR and cmMTB treated macrophages at day three. (**F**) Representative transmission electron microscopy images. Mitochondria are colored in red. (**G**) Quantification of the number of mitochondria per cell area. (**H**) Mitochondrial biomass assessed by flow cytometry using mitogreen staining at day three. Representative histograms (left) and dot plots showing the geomean fluorescence intensity (MFI), normalized to the cmCTR condition. (**I**) Superoxide production assessed by flow cytometry using mitoSox staining at day three. Representative histograms (left) and dot plots showing the MFI, normalized to the cmCTR condition. (**J**) Gene set enrichment plot of the OXPHOS geneset (hallmark collection of MSigDB). This plot shows the distribution of the genes of the oxidative phosphorylation pathway between macrophages exposed to cmCTR versus cmMTB for three days. (NES= 1.07; FDR= 0.583). Each circle within vertical plots represents a single donor. Mean +/- SD is shown. Statistical analysis: data with normal distribution; *, $p \leq 0.05$; **, $p \leq 0.01$; ***, $p \leq 0.001$; n.s. not significant.

**Supplemental Figure 2** (related to Figure 3)**. Modulation of aerobic glycolysis impacts HIV-1 infection of immunoregulatory macrophages.**

**(A-B)** Experimental design to evaluate the contribution of metabolic pathways to the exacerbation of HIV infection in cmMTB-treated cells. Briefly, monocytes from healthy subjects were differentiated in cmMTB for three days. At day two of differentiation, metabolic inhibitors were added to the culture medium. At day three, cells were infected by HIV-1 (NLAD8-VSVG) (for three further days) and HIV-1 infection of monocyte-derived macrophages (MDM) was measured. Monocytes from healthy subjects were treated with cmMTB or with TB-PE for three days. At day two of differentiation, metabolic inhibitors were added to the culture media. At day three, cells were infected by HIV-1 (NLAD8-VSVG) for three further days. (**C**) Flow cytometry analysis of cell viability, using LIVE/DEAD Violet Viability/Vitality Kit, at day three. (**D-G**) Analysis of MDM infection upon GSK 2837808A (GSK) treatment by microscopy. (**D**) Representative IF images: HIV-Gag (magenta) and nuclei (DAPI, grey). Scale bar, 100 µm. (**E**) Lactate release measured at day three (24 h upon treatment); quantification of MDM infection index (**F**, n= 9 donors) and MDM fusion index (G, n= 9 donors), normalized to the control condition (CTL= cmMTB w/o treatment). (**H-I**) Analysis of HIV-1 infection of cmMTB-treated macrophages upon drug treatment by flow cytometry. (**H**) Representative dot plots of HIV-Gag signals and gating strategy for selection of infected cells. (**I**) Quantification of % of infected cells (n= 8 donors), normalized to the control condition (CTL= cmMTB w/o treatment). Statistical analysis: data with normal distribution; *, p≤0.05; **, p≤0.01; ***, p≤0.001; ****, p≤0.0001.

**Supplemental Figure 3** (related to Figure 4)**. Glycolysis favors cell-to-cell transfer of HIV-1 and TNT formation in immunoregulatory macrophages.**
**(A)** Experimental setup used for the co-culture (also referred to as a transfer assay, Figure 4A-B). Briefly, monocytes from healthy subjects were differentiated in cmMTB for three days and cells were either infected by HIV-1 NLAD8-VSVG pseudo typed strain for additional three days (donor cells) or stained with CellTracker (recipient cells). At day six, donor cells were mixed with non-infected MDM stained with CellTracker (recipient cells) at a 1:1 ratio. Oxamate was added either at day two or 30 min after the coculture. The cells were fixed after 24 h of coculture and stained for HIV-1 Gag protein. HIV-1 transfer from infected to uninfected cells was determined at day 7. **(B-D)** Analysis of TNT formation. Monocytes from healthy subjects were treated with cmMTB for 3 days and, at day two of differentiation, UK5099, oxamate or GSK 2837808A (GSK) were added to the culture media. At day three, cells were infected by HIV-1 (NLAD8-VSVG) for three further days. (**B**) Representative confocal microscopy images: F-actin (grey, inverted). Red arrowheads show thick TNTs. Scale bar, 20 µm. (**C**) Quantification of the percentage of cells forming thick (defined as positive for both F-actin and microtubules (not shown), left) and thin (defined as positive for F-actin and not for microtubules, right) TNTs (n= 7-9 donors, at least 200 cells/condition/donor). (**D**) Quantification of the percentage of cells forming both types of TNTs (n= 8 donors, at least 200 cells/condition/donor). Mean +/- SD is shown. Statistical analysis: data with normal distribution; *, p≤0.05; **, p≤0.01; ***, p≤0.001.

### Supplemental Movie legend

**Supplemental Movie 1** (Related to Figure 4C).
Time-lapse of confocal microscopy images showing the fusion of HIV-1-infected (HIV-GFP viruses, grey) MDMs. One image every one min and 30 sec.

### Table 1

| Reagent type (species) or resource | Source or reference | Identifiers |
|---|---|---|
| **Critical Commercial Assays** | | |
| Lactate Kit | Wiener | Cat# 1999795 |
| 2-NBDG | Invitrogen | Cat# N13195 |
| MitoSOX Red Mitochondrial Superoxide Indicator | Invitrogen | Cat# M36008 |
| Cell tracker Green | Thermofisher Scientific | Cat# C7025 |
| Mitotracker DeepRed | Thermofisher Scientific | Cat# M22426 |
| CellROX Deep Red | Thermofisher Scientific | Cat# C10422 |
| Mouse anti-human CD14 microbeads | Miltenyi Biotec | Cat# 130-050-201 |
| LS magnetic columns | Miltenyi Biotec | Cat# 130-042-401 |
| Cell Dissociation Buffer | Thermo Fisher Scientific | Cat# 13151014 |
| Phalloidin AlexaFluor 488 | Thermo Fisher Scientific | Cat# A12379 |
| CellTracker Green CMFDA Dye | Thermo Fisher Scientific | Cat# C7025 |
| Fluorescence Mounting Medium | Agilent Technologies | Cat# S302380-2 |
| Ficoll-Paque Plus | Cytiva | Cat# 17144003 |
| Trypsin-EDTA (0.05%) | Gibco | Cat# 25300054 |
| Rpmi 1640 Medium, no glucose | Fisher Scientific | Cat# 11-879-020 |
| Glucose Solution | Thermo Scientific | Cat# A2494001 |
| **Bacterial and Virus Strains** | | |
| HIV-1 NLAD8-VSVG | Gift from Dr. S Benichou Institut Cochin, Paris, France | N/A |
| HIV-1 ADA Gag-iGFP-VSVG | Gift from Dr P. Benaroch Institut Pasteur, Paris, France | N/A |
| *M. tuberculosis* H37Rv | N/A | N/A |

| | | |
|---|---|---|
| **Biological Samples** | | |
| Buffy Coat from healthy donors | Etablissement Français du Sang, Toulouse, France | N/A |
| Patients-derived pleural effusions | Hospital F. J Muñiz (Buenos Aires, Argentina) | N/A |
| **Antibodies** | | |
| Anti-human HIF1-α | Cell Signaling | Cat# 36169S |
| LIVE/DEAD Fixable Aqua Dead Cell | Thermofisher Scientific | Cat# L34957 |
| Mouse monoclonal anti-HIV-1 p24 (clone KC57, FITC or RD1 coupled) | Beckman Coulter | Cat# 6604665/7 |
| Anti-human CD16 | Biolegend | Clone 3G8 |
| Anti-human CD163 | Biolegend | Clone GHI/61 |
| Goat anti-mouse IgG, AlexaFluor 488 | Thermo Fisher Scientific | Cat# A-10684 |
| Goat anti-Mouse IgG, AlexaFluor 555 | Cell Signaling Technology | Cat# 4409 |
| Anti-human CD169 | BioLegend | Clone 7-239 |
| **Chemicals, Peptides, and Recombinant Proteins** | | |
| Human M-CSF | Peprotech | Cat# 300-25 |
| TNTi | Pharmeks | N/A |
| Roswell Park Memorial Institute medium (RPMI 1640) | Gibco | Cat# 21875034 |
| Triton X-100<br>IF (0,3%)<br>WB (1%) | Sigma-Aldrich | Cat# T8532 |
| Tween 20 | Sigma-Aldrich | Cat# P9416 |
| UK5099 | MedChem Express | Cat# HY-15475 |
| GSK2837808A | MedChem Express | Cat# HY-100681 |
| Oxamate | Santa Cruz Biotechnology | Cat# sc-215880A |
| Fetal Bovine Serum (FBS) | Sigma-Aldrich | N/A |
| Ficoll-Paque Plus | Cytiva | Cat# 17144003 |
| Phosphate Buffer Saline (PBS) | Gibco | Cat# 14190144 |
| Bovine Serum Albumin | Euromedex | Cat# 04-100-812-C |
| Tris-buffered saline | Euromedex | Cat# 2-9134-10 |
| Paraformaldehyde | Delta Microscopies | Cat# D15714 |

| | | |
|---|---|---|
| Sucrose | Sigma-Aldrich | Cat# S0389 |
| Glutaraldehyde | Delta Microscopies | Cat# D16220 |
| **Software and Algorithms** | | |
| ImageJ | ImageJ | www.imagej.nih.gov/ij |
| Prism (v9) | GraphPad | www.graphpad.com |
| FACS DIVA | BD Bioscience | http://www.bdbiosciences.com/ |
| FlowJo v10 | FlowJo | https://www.flowjo.com/ |
| ZEN Black | Zeiss | https://www.zeiss.fr/microscopie/produits/microscope-software/zen.html |
